# Constant-pH MD Simulations of Lipids

**DOI:** 10.1101/2024.12.06.627182

**Authors:** Marius F.W. Trollmann, Paolo Rossetti, Rainer A. Böckmann

## Abstract

Constant pH Molecular Dynamics (CpHMD) simulations represent a cutting-edge computational approach for studying biological systems with remarkable realism. Recent advancements have enhanced the accessibility and efficiency of CpHMD, significantly reducing the performance overhead compared to traditional constant-protonation MD simulations. This chapter guides the reader through the application of CpHMD to investigate the pH-dependent behavior of Cationic Ionizable Lipids (CILs) — a critical component of Lipid Nanoparticles (LNPs), which are among the most promising platforms for drug delivery. LNPs, including those employed in mRNA-based vaccines, played a pivotal role in the global response to the SARS-CoV-2 pandemic, underscoring their potential in modern medicine. The chapter begins with a comprehensive introduction to the fundamental concepts of LNPs and provides a step-by-step protocol for setting up simulations of membranes containing CILs to calculate their apparent pK_a_. This parameter is crucial for governing the *in vivo* behavior of LNPs, where precise control is essential to optimize delivery efficiency while minimizing toxicity. By showcasing the ability of CpHMD simulations to unravel the intricate relationship between pH-dependent protonation, membrane structure, and lipid distribution, this chapter highlights their potential to inform the rational design of novel LNP formulations.

## 1 Introduction

### 1.1 pH and its importance for membranes

The concept of pH, introduced by S. P. L. Sørensen in 1904, provides a measure of the acidity or basicity of a solution — a critical parameter in biological systems. Living organisms tightly regulate cellular and extracellular pH through various mechanisms [55, 7], as hydrogen ion concentrations significantly affect the structure, dynamics, and functions of proteins, lipids, and nucleic acids [20, 63, 6, 3].

Titratable groups in molecules exhibit varying tendencies to (de)protonate, determined by their pK_a_ value, the pH at which protonated (*HA*) and deprotonated (*A*^−^) forms of the molecule are present in equal concentrations. The acid dissociation constant *K*_*a*_ relates to pK_a_ as:

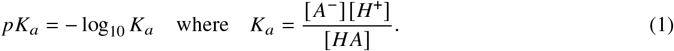

The pK_a_ of an isolated titratable group in aqueous solvent, termed the *intrinsic* pK_a_, is highly sensitive to its chemical environment. Both intramolecular and intermolecular interactions can alter the pK_a_ from its intrinsic value [64, 70].

The intramolecular environment involves the presence of electron-withdrawing and electron-donating groups, resonance effects [65], steric hindrance [40], and the formation of intra-molecular hydrogen bonds [39, 48]. Intermolecular factors include electrostatic interactions, hydrogen bonding, molecular crowding, solvation [47], and, related, the dielectric constant of the medium [44, 10]. For example, changes in solvent polarity affect the solvation energy of charged species, causing pK_a_ shifts [77, 58].

Titratable groups in membranes, proteins, and nucleic acids often display pK_a_ values that deviate from the intrinsic pK_a_ values of the respective isolated functional groups. These shifts, driven by the local environment, enable dynamic (de)protonation events under physiological conditions. Such behavior plays a critical role in cellular processes involving proteins [64, 35, 30, 4, 54], membranes [19, 53], and nucleic acids [74].

This chapter will focus on the application of constant-pH molecular dynamics (CpHMD) simulations to titratable lipids and their role in the study of membrane structure and dynamics. Many membrane lipids, both natural and synthetic, contain titratable groups, such as the phosphate group in phospholipids. Lipids with pK_a_ values within the physiological pH range (2–8.5) are of particular interest due to their protonation dynamics in various cellular environments. Several drugs and formulations leverage pH variations across the body to enhance the delivery of active ingredients [5]. Lipid nanoparticles (LNPs) incorporating cationic ionizable aminolipids (CILs) are especially noteworthy due to their pronounced pH-dependent behavior [11, 70]. A

### 1.2 Ionizable lipids in drug delivery

Cationic aminolipids, the precursors of ionizable aminolipids, feature a permanently charged (non-titratable) trimethylammonium group. These cationic lipids effectively bind to negatively charged nucleic acids, improving encapsulation efficiency and facilitating cellular delivery [23]. However, the permanent positive charge of the trimethylammonium group leads to several limitations, including poor pharmacokinetics (PK) and *in vivo* toxicity. The constant surface charge results in non-specific membrane fusion, causing hemolysis [33], and triggers pro-apoptotic pathways upon LNP endocytosis [15]. Moreover, cationic liposomes are rapidly cleared from circulation due to interactions with plasma proteins and charge-sensitive cellular uptake [62].

To overcome these challenges, CILs containing a tertiary amine group were developed. These lipids exhibit tunable charge properties that improve their safety and efficacy profiles. Notably, three ionizable aminolipids — DLin-MC3-DMA, ALC-0315, and SM-102 — have been approved for clinical use [24]. The *intrinsic* pK_*a*_ of isolated, solvated CILs typically ranges from 8.5 to 9.5 [11], while their *apparent* pK_*a*_ within LNPs shifts to 6-7. This shift enables CILs to remain predominantly neutral in the bloodstream (pH 7.4) and protonated within the acidic endosomal lumen (pH 6) [51, 28] (see Figure 1A). CILs offer the advantage of being predominantly neutral in the blood circulation (pH 7.4) while being prevalently protonated in the endosomal lumen (pH 6) [51, 28] (see scheme in Figure 1A). The reduction in surface charge at neutral pH minimizes non-specific interactions with tissues and proteins, such as uptake by hepatocytes [37], thereby improving the safety profile and pharmacological activity of LNPs [61].

**Fig. 1:**
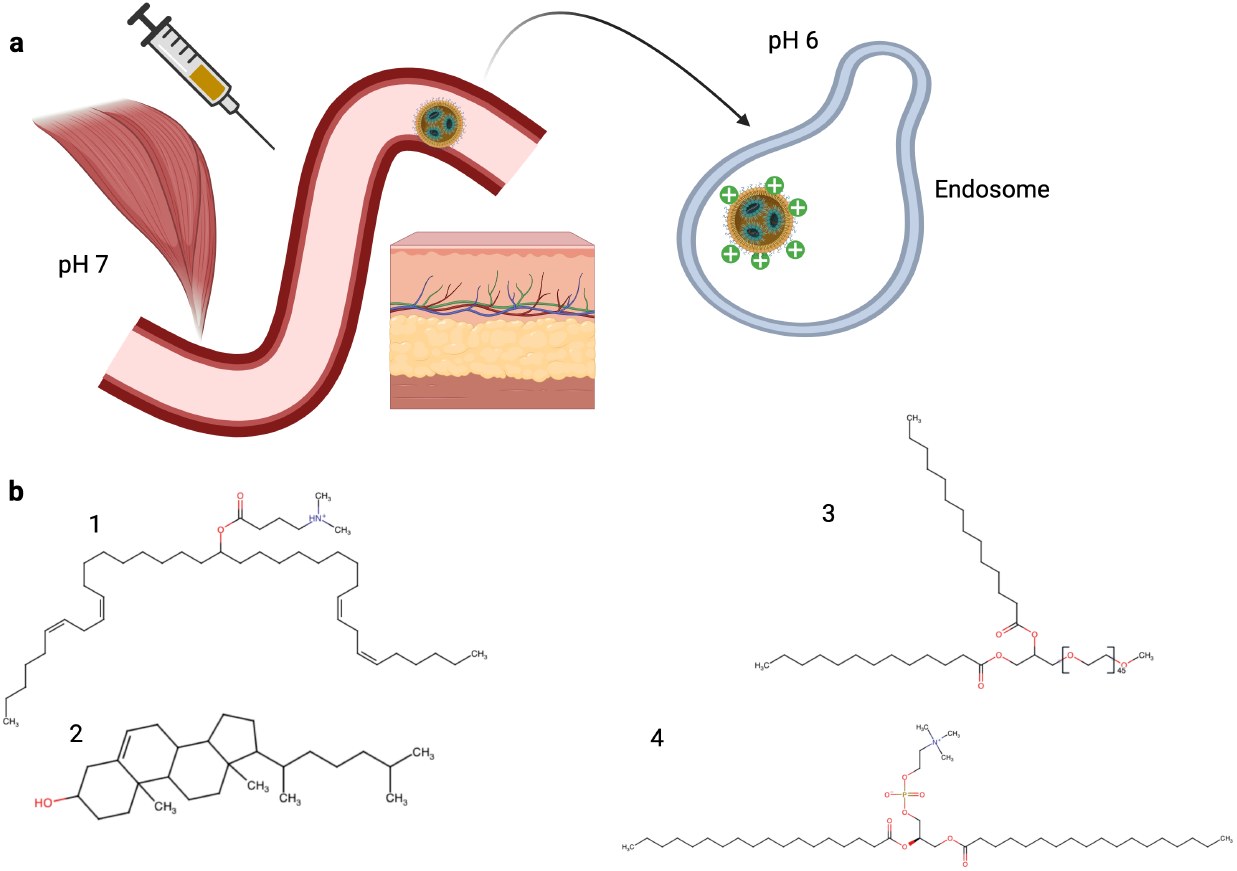
(**A**) LNPs can be administered intravenously, subcutaneously or intramuscularly. At near-neutral pH in these environments, the LNP surface carries minimal surface charge. However, the charge increases inside the more acidic endosomal lumen. (**B**) Composition of a typical LNP: (1) the ionizable aminolipid D-Lin-MC3-DMA, (2) cholesterol for structural stability, (3) PEGylated lipid (DMG-PEG 2000) to enhance circulation time and reduce aggregation, and (4) the helper phospholipid DSPC, which aids in forming a stable LNP surrounding lipid monolayer.

CILs are highly customizable, with both their intrinsic pK_a_ (modulated by the electron density on the nitrogen atom) and apparent pK_a_ (influenced by LNP composition) being adjustable to meet specific needs. Additional properties, including molecular shape and biodegradability, are also critical in CIL design. The conical shape of most CILs enhances membrane fluidity [69], promoting endosomal escape [24]. Biodegradability, which influences tolerance and immunogenicity, can be modulated by incorporating metabolically labile groups, such as esters with varying degrees of steric hindrance [51, 46, 13].

In clinical formulations, CILs are combined with three other key LNP components: helper phospholipids, cholesterol, and PEGylated lipids [16] (see Figure 1B). These formulations are tailored to the specific payload, as nucleic acid loads vary in size and structural organization [28]. Among the factors influencing LNP performance, the apparent pK_a_ and CIL shape have emerged as primary predictors of *in vivo* activity [73]. Consequently, structural insights can guide the rational design of LNPs with enhanced safety and efficacy.

### 1.3 The relationship between intrinsic and apparent pK_a_

The pK_a_ of a titratable group is highly influenced by its environment, including factors such as the dielectric constant of the surroundings, molecular interactions, molecular flexibility, and the ionic strength of the solvent [51, 59, 10]. As a result, the apparent pK_a_ of titratable groups can shift significantly when transitioning from an aqueous environment to the hydrophobic core of a membrane or lipid nanoparticle.

In the heterogeneous LNP environment, the apparent pK_a_ of cationic ionizable lipids (CILs) is highly dependent on their location within the nanoparticle [70]. For example, CILs complexed with nucleic acids in the LNP core form salt bridges with phosphate groups, leading to an increase in the pK_a_ of the tertiary amine group [70]. Also, nucleic acid phosphate groups, which typically have very low intrinsic pK_a_ values [67], can experience pK_a_ shifts due to structural changes like RNA folding [74]. In the hydrophobic, low-dielectric core of the LNP, most aminolipids remain neutral. Conversely, CILs at the LNP surface are exposed to the aqueous phase and interact with helper lipids’ phosphate groups, which increases their protonation and apparent pK_a_. However, shielding by the polyethylene glycol (PEG) groups of the PEGylated lipids can reduce solvent exposure in some surface regions [69], stabilizing the neutral form of aminolipids and lowering their pK_a_. Unlike the intrinsic pK_a_, the apparent pK_a_ is less precisely defined. In the context of LNPs, it reflects the aggregate protonation behavior of the lipid mixture. Experimental studies indicate that the apparent pK_a_ of LNPs is typically about 3 pK units lower than the intrinsic pK_a_ of isolated aminolipids [51, 11], indicating that lipid-lipid interactions reduce the basicity of CILs. The apparent pK_a_ is also formulation-dependent; changes in the molar ratios of LNP components can lead to significant shifts [51, 76, 66]. Additionally, the method of measurement affects the reported apparent pK_a_, which should be regarded as a property of the LNP formulation rather than of individual lipids.

Optimizing the apparent pK_a_ of LNPs is critical for improving their performance in drug delivery. High-throughput computational methods [29] are accelerating the design of new CILs and formulations, enabling rapid exploration of potential candidates. Current studies suggest that the optimal apparent pK_a_ range for promoting endosomal escape lies between 6 and 7. Small variations in pK_a_ can significantly influence organ tropism and site-specific delivery [51, 37, 31, 66]. These effects are closely tied to changes in the protein corona, the dynamic layer of proteins associated with the LNP surface [17, 25, 52].

The following section delves into the experimental and computational techniques used to study lipid nanoparticles. Special emphasis is placed on constant-pH molecular dynamics (CpHMD) simulations as a versatile tool for probing pH-responsive LNP behavior at atomistic resolution.

### 1.4 Methods for studying aminolipids and lipid nanoparticles

The study of LNPs and their components, such as aminolipids, requires a multidisciplinary approach encompassing four key categories: physicochemical experiments, biological assays, *in vivo* studies, and computational techniques.

Physicochemical methods such as dynamic light scattering (DLS), electron microscopy (EM), and small-angle X-ray scattering (SAXS) are essential for characterizing LNP structure and organization. For example, EM has revealed critical details about mRNA-loaded LNPs, including water- and RNA-rich electron-lucent regions surrounded by an electron-dense amorphous core and a surface layer [11]. SAXS studies further demonstrated the self-assembly of LNPs at acidic pH in the presence of nucleic acids [75]. Additionally, *ζ*-potential measurements at varying pH values and the apparent pK_a_ determined through TNS assays (using the fluorogenic probe 6-(ptoluidino)-2-naphthalenesulfonic acid sodium salt) are critical for understanding the equilibrium between charged and neutral aminolipids, which directly affects LNP functionality.

Cellular assays are indispensable in understanding the complex factors that influence LNP delivery efficiency and toxicity. Key methods include caspase activation assays [15], immune cell activation assays [22], and protein expression assays, which collectively assess safety, activity, and immunogenicity—fundamental parameters for evaluating LNP performance [28].

Physicochemical and cellular assays, while valuable, cannot fully capture the interactions of LNPs with various biomolecules and bodily tissues. Consequently, *in vivo* studies are essential for capturing the full scope of ADMET (absorption, distribution, metabolism, excretion, and toxicity) parameters, such as organ tropism [57], hepatotoxicity [24], and biodegradability [46]. Discrepancies between *in vitro* and *in vivo* outcomes [22] often arise due to protein corona formation and individual variability in protein and receptor expression levels. The composition of the protein corona is primarily dictated by LNP surface properties [14] but is also influenced by inter-individual differences [57, 45]. Ethical and promising methods, such as the use of peripheral blood mononuclear cells (PBMCs) from human donors, are valuable for studying these effects [57, 45].

Molecular dynamics (MD) simulations provide an atomistic view on LNP structure and behavior. Both all-atom and coarse-grained (CG) models have been used to explore experimentally characterized LNP formulations [69, 42]. Whole-LNP simulations are computationally intensive, but an alternative approach involves simulating cuboid sections of the LNP with consistent surface area-to-volume ratios, significantly reducing computational costs [49, 69]. These simulations enable e.g. the study of phenomena such as LNP fusion with endosomal membranes [42].

Molecular dynamics (MD) studies to date have predominantly employed classical constant-protonation MD simulations to explore how varying ratios of protonated and neutral species influence the surface properties of LNPs [69, 19, 49, 34]. These studies have shown that neutral aminolipids preferentially partition into the hydrophobic core of the LNP due to their higher hydrophobicity and conical, non-bilayer-forming shape [69, 19]. In contrast, protonated aminolipids, characterized by increased polarity and larger effective headgroup sizes, tend to organize into bilayers [75].

While these computational studies have primarily relied on classical MD simulations with constant protonation, recent advancements have made constant-pH MD (CpHMD) simulations more accessible. These developments have reduced computational costs and simplified the setup process for both all-atom and CG simulations [1, 36, 27, 12, 68, 8]. Consequently, CpHMD is poised to become a widely adopted approach for studying biologically relevant processes that depend on pH variations.

Most CpHMD approaches utilize *λ*-dynamics [43], where a one-dimensional *λ*-coordinate is used to define the protonation state of a residue [1]. The *λ*-coordinate is influenced by three critical factors: (1) the user-specified pH, (2) the reference pK_a_ values of the respective titratable groups, and (3) electrostatic interactions with surrounding atoms [1]. Among these, the third factor is particularly significant, as it allows CpHMD to accurately capture the effects of the local environment on protonation states. The first two factors highlight the importance of experimental data in ensuring the reliability of simulations. Since CpHMD only allows for accurately calculating differences in the energy of the protonation states of a titratable group, providing an accurate reference intrinsic pK_a_ is essential for replicating pH-dependent behaviors observed experimentally.

This chapter will demonstrate how CpHMD can be used to derive titration curves for aminolipids in both aqueous and membrane environments, enabling the determination of apparent pK_a_ values. It provides a detailed protocol including the parameterization of the aminolipid MC3 for CpHMD, as well as an application study. This protocol is based on procedures outlined in previous works for setting up CpHMD simulations of proteins [1, 9, 36]. A significant difference from those studies is that no modifications to dihedral potentials were made, as it was done for the CHARMM36m protein force field [1, 9].

## 2 Materials

The workflow outlined in this chapter involves using several software tools that are freely available for academic purposes, though some may require registration. The provided command-line tools and software are mostly compatible only with UNIX-based systems (i.e., Linux or MacOS), so a basic understanding of these systems is assumed. The following programs need to be installed:

- `GROMACS` with the constant-pH implementation by Aho *et al*. [1]: A versatile software package for MD simulations with a high performance on CPU as well as GPU nodes. The constant-pH extension was implemented in GROMACS 2021.4 and is available under https://gitlab.com/gromacs-constantph/constantph (last accessed 06.10.2024)
- `phbuilder` [36]: A collection of Python-based command-line tools to setup constant-pH simulations. Installable via the pip-package manager for Python (Documentation under https://gitlab.com/gromacs-constantph/constantph, last accessed 06.10.2024).
- `PyMOL` [60]: Visualization of molecular structures and molecular dynamics trajectories. PyMOL is available at the offical Schroedinger website (https://www.pymol.org, last accessed 07.10.2024) (paid or educational license-required). An offical GitHub-repository containing the open-source version of `PyMOL` is also available under https://github.com/schrodinger/pymol-open-source (last accessed 07.10.2024).

Additional scripts are required to facilitate the setup of the thermodynamic integration and the titration simulations. All three scripts can be found in the `phbuilder` GitLab repository https://gitlab.com/gromacs-constantph/phbuilder (Last accessed: 10.10.2024):

- `create parameterization.py`: Modifies .mdp files and prepares .tpr files for the thermodynamic integration employed to estimate 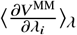.
- `fit parameterization.py`: Performs a fitting routine to estimate the polynomial coefficients for the gradient of 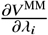 along the *λ*-coordinate.
- `create titration.py`: Modifies .mdp files and prepares .tpr files for multiple independent replica simulations at different pH levels to obtain a titration curve from simulations.

Furthermore, we recommend to register for a CHARMM-GUI account (https://charmm-gui.org, last accessed 06.10.2024) to obtain the parameter files required for the protocol and to facilitate the setup of complex membrane systems.

Beside the computational tools, another crucial parameter for the setup of constant-pH simulations are the intrinsic pK_a_ values of the titratable groups present in the simulation system. Note that only the difference between pH and pK_a_ is used within constant-pH MD, since the difference in Gibbs free enthalpy between protonated and deprotonated states is given by

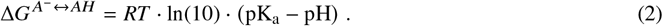

The intrinsic pK_a_ value must be provided to allow for a quantitative comparison between titration curves obtained from simulations and experiments. For example, a suitable reference value would be the pK_a_ of the titratable molecule in an aqueous medium (e.g., water with a physiological salt concentration of 150 mM Na^+^Cl^-^) at an “infinite “ dilution, referred to here as the intrinsic pK_a_. For hydrophobic molecules, like aminolipids, this value can be hardly obtained from experiments due to their low critical micellar concentration (cmc). The intrinsic pK_a_ of such molecules is typically determined either by using water-soluble analogs or from theoretical modeling approaches.

For example, Carrasco *et al*. [11] employed both a software package from Advanced Chemistry Development, Inc. (ACD/Labs), and NMR spectroscopy on water-soluble analogs of various ionizable lipids to estimate their intrinsic pK_a_. Differently, Hamilton *et al*. [29] used the machine-learning-based software suite Epik [38] to predict intrinsic pK_a_ values for four different ionizable lipids. Both studies reported shifted pK_a_ values in lipid mixtures, such as LNPs, referred to here as the *apparent* pK_a_. This observation will be discussed in the next section in comparison with results of the CpHMD simulations performed in this protocol.

## 3 Methods

This section provides a step-by-step guide, detailing the process from the parameterization of an aminolipid to the simulation of a complex model membrane under constant-pH conditions. The workflow is designed for UNIX-based systems (e.g., Linux or macOS); however, syntax and commands may vary depending on the specific operating system, command-line tools, or software versions used. Readers are encouraged to verify and adapt each step to ensure accuracy. The length and number of simulations described here are tailored to the requirements of the example lipid system and may need adjustment for different lipid types, membrane compositions, or target observables.

The aminolipid D-Lin-MC3-DMA (MC3) is used as the model system in this chapter due to its relatively simple structure and its widespread application in both scientific research and therapeutic formulations. A prominent example of its use is in the LNP-based siRNA therapeutic Onpattro [2]. Additionally, molecular mechanics parameters for both protonated and deprotonated states of MC3, compatible with the CHARMM36 force field, are already available [50].

### Important Note

This tutorial utilizes default MD parameters provided by CHARMM-GUI and `phbuilder`. However, specific adjustments have been made for simulations involving thermodynamic integration for parameter estimation, and initial validation runs (e.g., unbiased simulations and titration). These adjustments include adopting more conservative settings for neighbor list generation, as recommended by Kim *et al*. [41].

Specifically, the `rlist` value was fixed at 1.35 nm by disabling the automated buffer setting in `GROMACS` (`verlet-buffer-tolerance=-1`). Neighbor searching was conducted every 20 steps (`nstlist = 20`), coinciding with the coupling intervals for both temperature (`nsttcouple = 20`) and pressure (`nstpcouple = 20`). A repository with all the parameters and structures will be made available to Zenodo, after the final acceptance of the manuscript.

### Protocol

1. Start with preparing a membrane patch containing the ionizable lipid using the `Membrane Builder → Bilayer Builder` provided by CHARMM-GUI.
  - Select the `Membrane Only System` option to start the system building process.
  - Choose a `Hydration number` of 50 water molecules per lipid molecule and select the lateral length of the simulation box to be based on the `Number of lipid components`.
  - For the membrane composition, select `72 DOPC`, `24 Cholesterol`, and `24 DLMC3H` (protonated MC3 with 18:2/18:2 acyl chains) for each leaflet. Note that the membrane composition includes all lipids typically used in LNP formulations, except for PEGylated lipids. Fractions of cholesterol and MC3 are reduced to 20 % each to prevent significant (and slow) lipid reorganization due to pH changes [69].
  - For `Step 3` of the `Membrane Builder`, ensure that the `Include Ions` box is checked. No or only a low salt concentration should be added to the simulations of the complex membrane composition, except for counter ions to ensure charge neutrality of the system. **Important note**: A physiological salt concentration is required when parameterizing the MC3 lipid for the constant-pH MD simulation (detailed below). Unfortunately, the CHARMM-GUI output contains only the force field parameters for atoms and molecules present in the prepared system. To simplify the setup of the parameterization simulations later, add a low Na^+^Cl^-^ concentration (e.g., 0.01 M) along with the counter ions. The few excess ions can either be removed from the system later or kept without significantly prolonging the equilibration process.
  - In the following steps, CHARMM-GUI assembles the requested system.
  - In `step 5`, select `CHARMM36m` as the required force field. Under `Input Generation Options`, choose `GROMACS` as an additional output format, and under `Equilibration Options` set the temperature to 310 K (physiological temperature).
  - Finally, in `step 6`, the generated structure along with the requested topologies are prepared for download.
2. The initial structure of the assembled membrane is prone to numerical instabilities, e.g., caused by overlapping atoms leading to high potential energies. For the initial equilibration, follow the workflow suggested by the CHARMM-GUI, as detailed in the executable `README` file. For the membrane simulation in this tutorial, only counter ions (i.e., 48 chloride anions) are added to the system. If a salt concentration was previously included to obtain the force field parameters for the sodium cation, the excess Na^+^Cl^-^ can be removed from the input files `step5 input.gro` and `topol.top` using a plain text editor such as vim or gedit. The default workflow involves multiple short equilibration simulations with position restraints on the lipids, followed by 10 independent “production “ simulations using the `step7 production.mdp` file, each with a duration of 1 ns. For this tutorial, only the final structure of one “production “ simulation is needed. The number of simulations can be adjusted in the `README` file before running the full equilibration workflow with: ~~~
csh README
~~~ The final equilibrated membrane structure, with all MC3 lipids protonated, corresponds to a highly acidic environment, with a pH much below the *apparent* pK_a_ of the titratable lipid within the lipid mixture.
3. The implementation of the constant-pH method by Aho *et al*. [1] uses three *λ*-dependent potentials that are added to the total potential energy function: (1) **V**^**bias**^(*λ*) ensures a proper sampling of the physical relevant states, i.e, *λ*≈ 0 (corresponds to aminolipid protonated) or *λ*≈ 1 (aminolipid deprotonated), during the simulation. (2) **V**^**pH**^(*λ*) induces the effect of the chosen pH on the protonation state of the titratable sites via an implicit proton bath. (3) **V**^**MM**^(*λ*) corrects the force field’s proton affinity and is defined as the negative of the deprotonation free energy for a titratable site *i*,

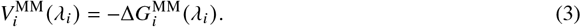

In contrast to *V*^bias^(*λ*) and *V*^pH^(*λ*), no analytical representation for *V*^MM^(*λ*) exists. To estimate *V*^MM^(*λ*_*i*_) for a titratable residue *i*, its gradient along the *λ*_*i*_-coordinate, i.e., 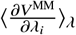, must be derived at the force field level. Following the approach of Aho *et al*. [1], a high-order polynomial is then fitted to the values of the gradient to enable an efficient and accurate calculation of the force contribution. There are two strategies to obtain the polynomial coefficients: running long thermodynamic integrations until the estimates for 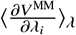 converge [1, 9], or using the two-step scheme presented by Jansen *et al*. [36]. In the two-step scheme, a “quick’n’dirty” thermodynamic integration is first performed (e.g., ≈1 ns) to obtain an initial guess for the polynomial coefficients. Followed by long unbiased sampling simulations (with the biasing barrier of *V*^bias^ (*λ*_*i*_) set to 0) to determine the (non-ideal) distribution of *λ*_*i*_. Boltzmann inversion is then employed to re-weight *V*^MM^ (*λ*_*i*_) with Δ*V*^MM^ = −*RT*· ln (*p* (*λ*_*i*_)) [36]. In this tutorial, the former approach of long thermodynamic integration is employed to obtain the polynomial coefficients. For that, the residue is typically placed in a water-filled box along with a buffer particle that compensates for changes in the charge of the titratable site to maintain a constant net charge within the box [1, 9] (see Fig. 2). The *λ*-coordinates are fixed to values between [−0.1, 1.1] with the constraint that *λ*_*BUF*_ + *λ*_*i*_ = 1.0. Keeping both *λ*-coordinates fixed during the simulation allows for the estimation of 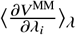.
  - Since only one molecule is required for the thermodynamic integration, extract a single MC3 lipid from the membrane created in **Step 1**. To do this with `GROMACS`, create an index (.ndx) file that contains an index group with just one MC3 residue using the `make ndx` command: ~~~
gmx make_ndx -f step5_input.pdb -o mc3_r1.ndx
~~~ If the first residue in `step5 input.pdb` is an MC3 lipid, you can create the index group with the command `ri 1`.
  - The structure is then extracted from the .gro file with the following command: ~~~
gmx editconf -f step5_input.pdb -n mc3_r1.ndx -ndef -o mc3_init.pdb -label A,
~~~ selecting the `r 1` group for the output.
  - The lipid is subsequently placed in a cubic box with a volume of 6×6×6 nm^3^ ~~~
gmx editconf -f mc3_init.pdb -o mc3_box.pdb -box 6 6 6
~~~
  - To facilitate the placement of the buffer atom in a later step, the nitrogen atom of the lipid is centered to the middle of the box. A corresponding index file is created with, ~~~
gmx make_ndx -f mc3_box.pdb -o center.ndx
~~~ selecting the nitrogen atom with `a N1`.
  - The `gmx trjconv` routine is used to center the nitrogen atom, ~~~
gmx trjconv -f mc3_box.pdb -s mc3_box.pdb -n center.ndx
            -center -o mc3_box_center.pdb
~~~ First, select the index group containing only the nitrogen (`N1`), followed by the index group for the lipid (`System`).
  - The TIP3 water model is applied in all subsequent simulations, so the box is filled up with a 3-point water model using, ~~~
gmx solvate -cp mc3_box_center.pdb -o mc3_solvated.pdb
~~~ **Important note**: The naming conventions for water molecules and their atoms differ between the output of `gmx solvate` (where the water is labeled as **SOL**) and the force field provided by CHARMMGUI (where water is labeled as **TIP3**). To resolve this discrepancy, the naming conventions from the `GROMACS` output are adopted in the `TIP3.itp` file.
  - The buffer particle is a single atom, comparable to an ion, and can be placed inside the system with, ~~~
gmx insert-molecules -ci BUF.gro -f mc3_solvated.pdb -o mc3_solvated_buf.pdb
 -ip positions_BUF.dat -replace SOL
~~~ `BUF.gro` contains a single buffer particle initially positioned at `0.0 0.0 0.0`, and `positions BUF.dat` specifies the coordinates for placing this particle in the system, which is also set to `0.0 0.0 0.0` (see Fig. 2).
  - After adding the buffer particle, a topology file should be created that includes the path to the force field files and a list of molecules present in the system: 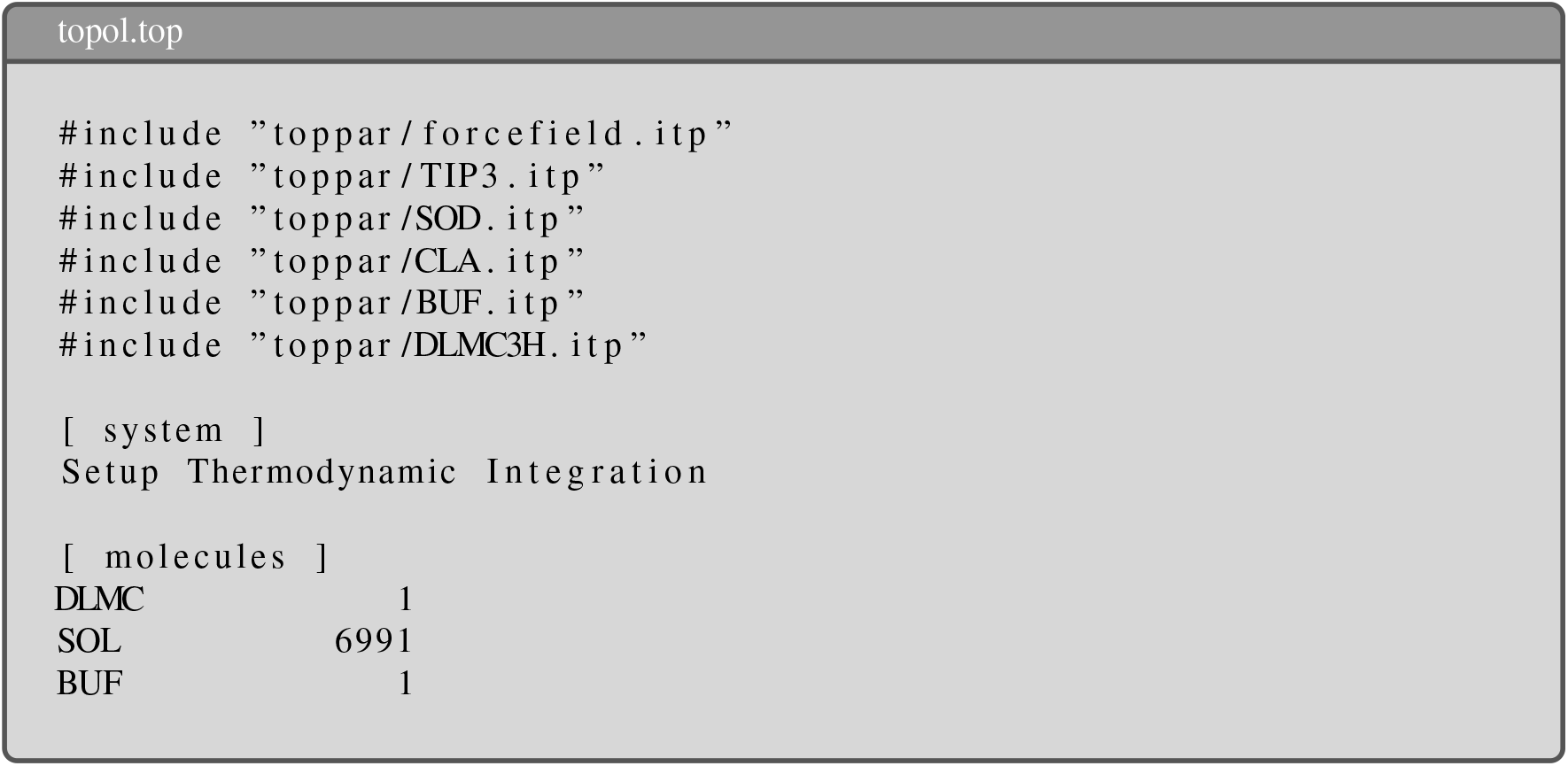
  - It is recommended to fix the positions of both the residue and the buffer particle during the simulations to minimize their interactions. This helps prevent spoilage in the estimate of 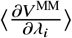.An example of the topology for the buffer particle together with appropriate position restraints [36] is given in the next box. 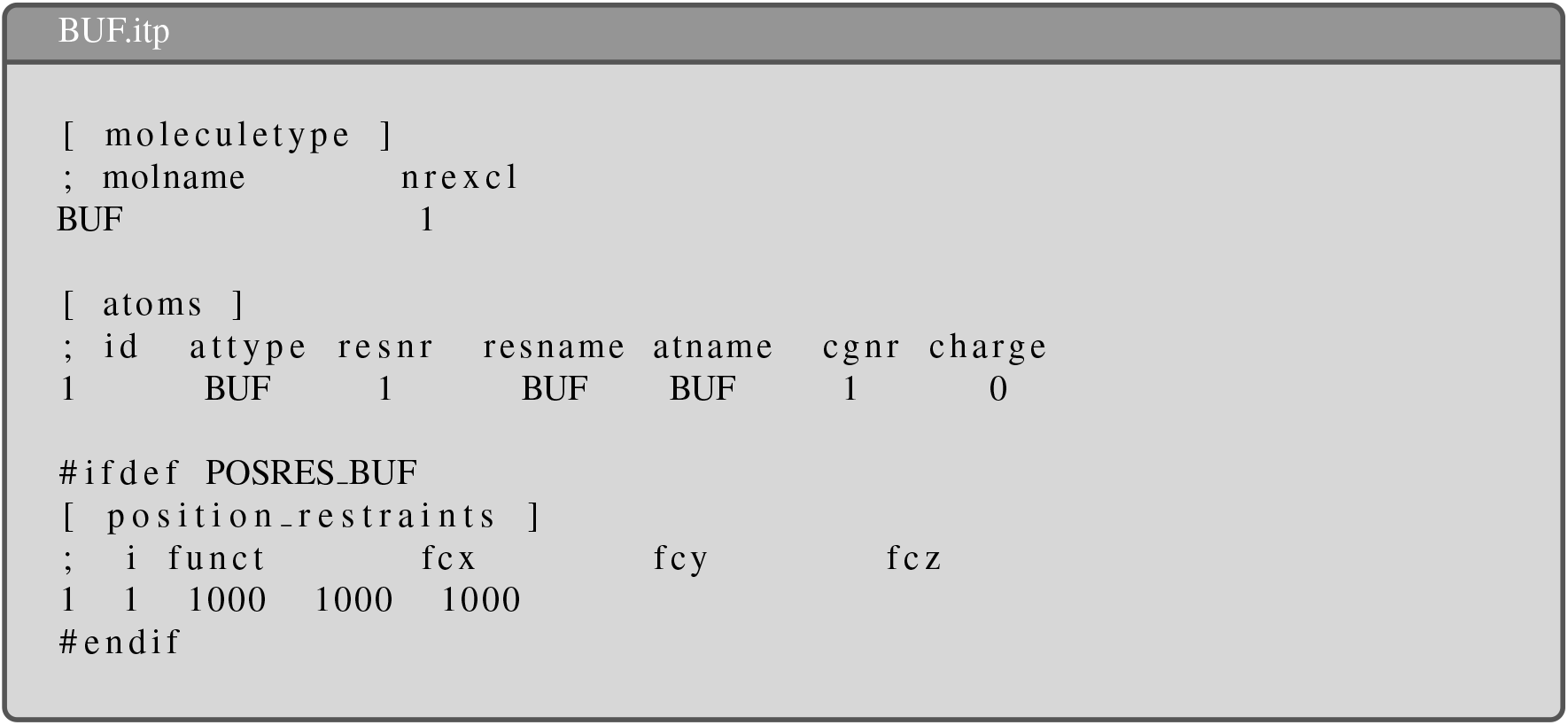 Since, the topology of the buffer particle introduces a new atom type `BUF`, the following lines must be added to the `forcefield.itp` under the respective directives: 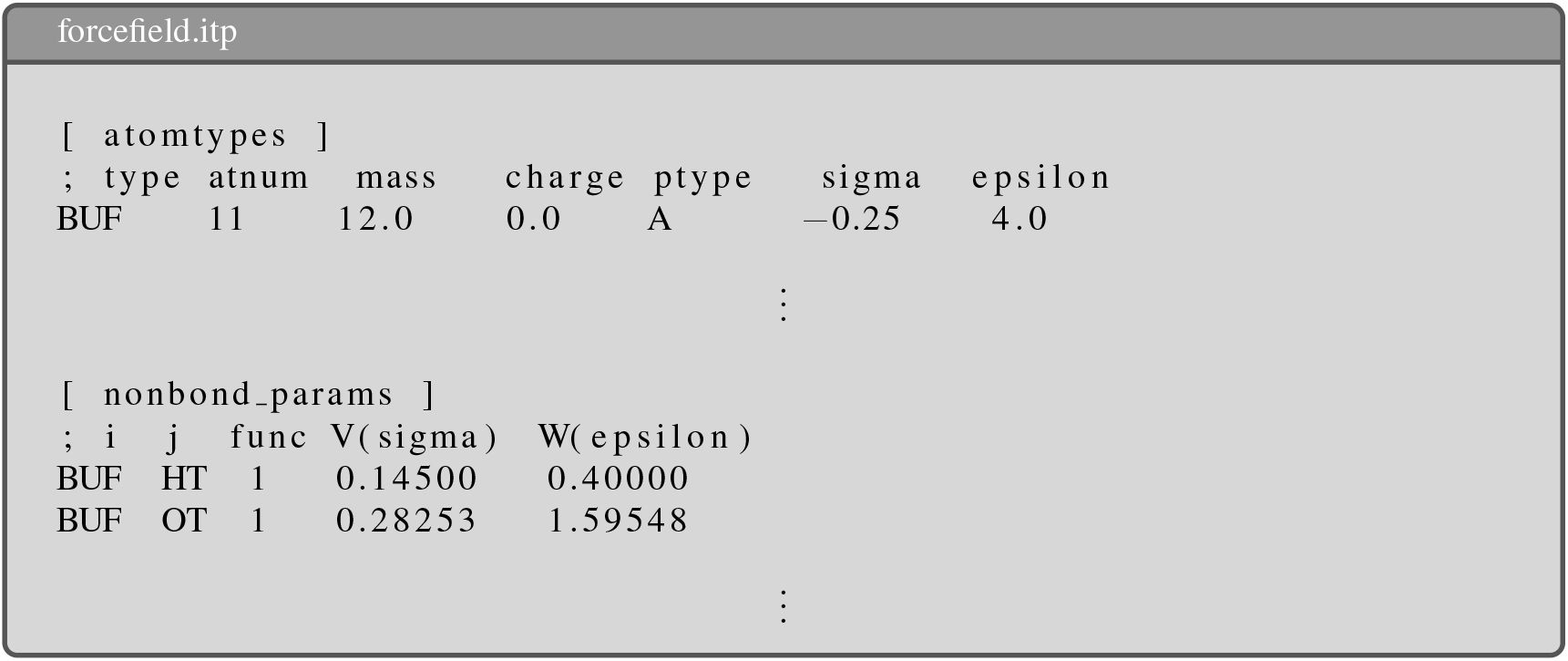 More information regarding the parameterization of the buffer particles is provided in Buslaev *et al*. [9].
  - The system setup is now almost complete. For the final steps, the Python-package `phbuilder` [36] is employed to generate the input files (e.g., .mdp and .ndx) for the constant-pH MD simulations, and to add ions to the system. A key input file for `phbuilder` is `lambdagrouptypes.dat`, which contains information such as the location of the `GROMACS` installation directory, the name of the force field directory, and the name of the atoms in the titratable sites. **Important note:** The order of the atoms should correspond to their atom indices as listed in the .itp file. 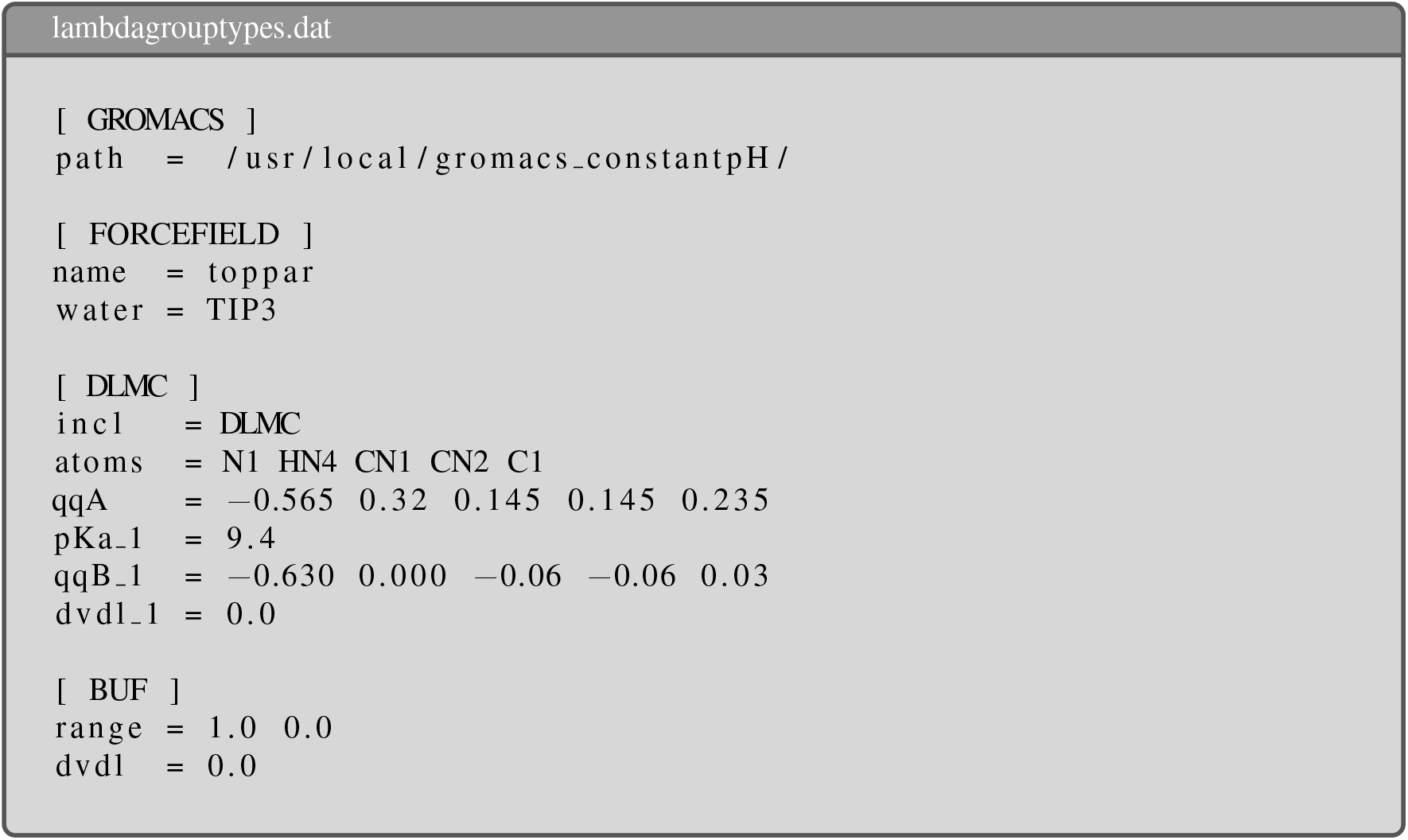 The `[DLMC]` section specifies the name of the MC3 lipid in the force field, the atoms found in the titratable site, the partial charges for both the protonated and deprotonated states [50], and the intrinsic pK_a_ of the aminolipid [11, 29]. To ensure that `phbuilder` correctly recognizes the titratable residue, the residue name of the aminolipid should be changed to `DLMC` in both `mc3 solvated buf.pdb` and `DLMC3H.itp`. Since the polynomial coefficients (`dvdl 1`) for the *V*^MM^(*λ*_*i*_) potential are unknown at this step, they are set to 0. During thermodynamic integration, the buffer particle (`[BUF]`) can take a charge between 0.0 and 1.0. Since the value of the *λ*_*BUF*_-coordinate is fixed, no polynomial coefficients for V^MM^(*λ*_*BUF*_) are required. The `neutralize` tool of `phbuilder` is employed at this step to add an ion concentration of 150 mM Na^+^Cl^-^ to the system: ~~~
phbuilder neutralize -f mc3_solvated_buf.pdb -conc 0.15 -pname SOD -nname CLA
                     -rmin 1.0 -nbufs 0 -ignw -v
~~~ This command adds 150 mM Na^+^Cl^-^ to the system, based on the current solvent volume. The residue names follow the CHARMM-GUI naming conventions, with `SOD` representing the sodium ion and `CLA` representing the chloride ion. It is generally not recommended to perform simulations with a non-zero net charge when using the Particle-Mesh-Ewald (PME) algorithm [18], as it can lead to inaccuracies in long-range electrostatics calculations [32]. Therefore, a chloride anion is added to neutralize the (combined) charge of the aminolipid and the buffer particle. The flag `-rmin` ensures that particles are placed at least 1.0 nm away from the lipid. The `-ignw` flag ignores all warnings raised by `grompp`; this is generally **not** recommended but is necessary here to bypass issues with `grompp` correctly reading longer atom names (e.g., `H218R`). The output structure is stored in a .pdb format (`phneutral.pdb`).
  - After the system is finally prepared, the .mdp parameters for the simulations can be created with the `genparams` subroutine. To initially protonate the MC3 lipid, a `phrecord.dat` file is defined, which is automatically read by `genparams`. This file contains the name of the lipid as specified in the force field, its residue number, the chain label (which is empty in this case), and the initial value of the *λ*-coordinate. 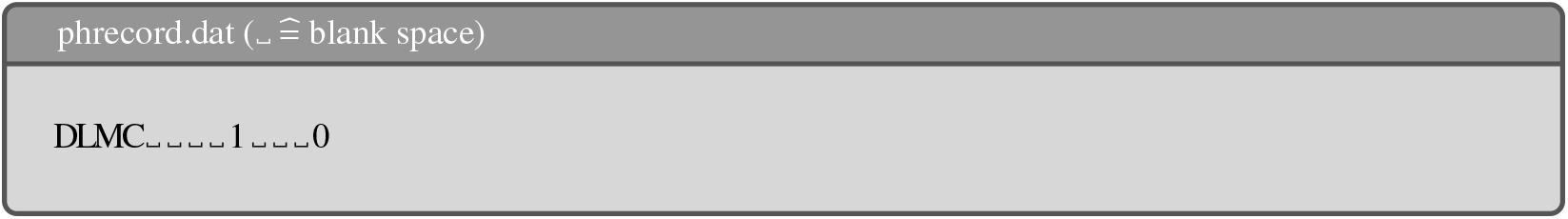 ~~~
phbuilder genparams -f phneutral.pdb -cal -dwpE 0.0 -ph 9.4 -v
~~~ The `-cal` flag sets `lambda-dynamics-calibration = yes` in the `MD.mdp` file, which fixes the *λ*-coordinates during the simulations. Setting `-dwpE` to 0.0 removes the barrier height imposed by the bias potential *V*^bias^ (*λ*_*i*_) [1], and choosing the pH equal to the pK_a_ eliminates the contribution of *V*^pH^ (*λ*_*i*_). However, both potentials are ignored anyways due to `lambda-dynamics-calibration = yes`. Further, `genparams` creates an index file containing the groups `LAMBDA1` and `LAMBDA2`, which are used in the .mdp file to define the group of atoms for the partial charge scaling for each titratable site. Note that `genparams` sets the reference temperature to 300 K by default. If a different value (e.g., 310 K) is required, it must be set manually. **Important note**: The position restraint for the `N1` atom in the aminolipid, as defined in `DLMC3H.itp`, is only applied in the z-direction. However, the parameterization requires restraints in all directions. Therefore, the `[position restraints]` directive must be modified accordingly, as shown in the next box. 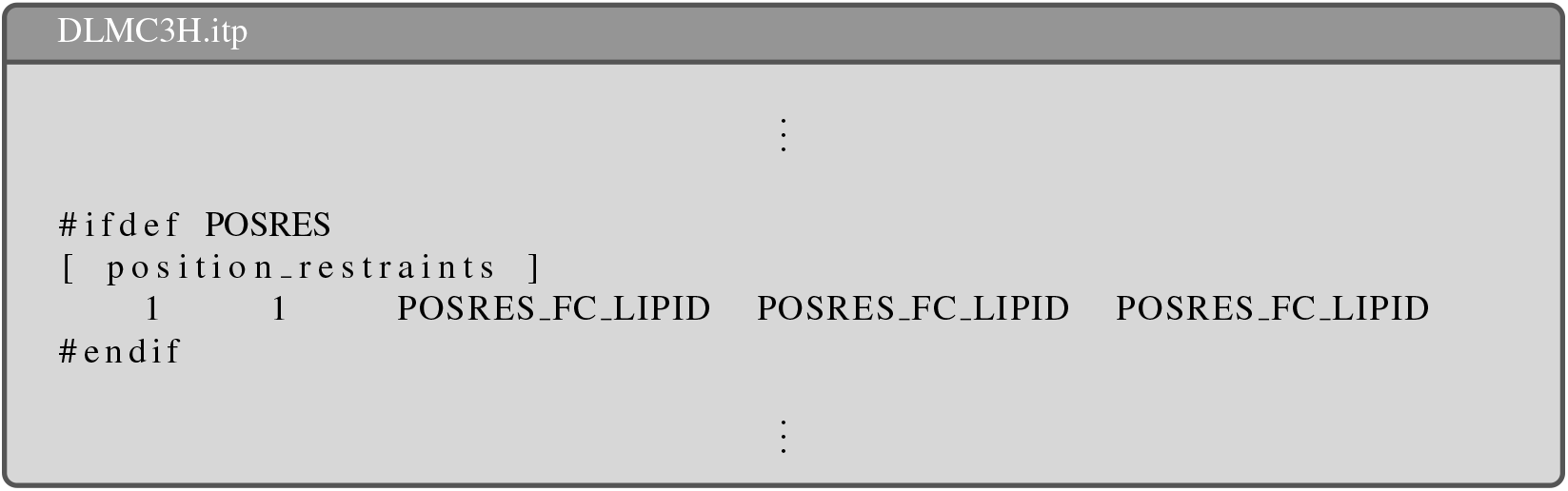 The magnitude of the restraint force for the aminolipid is then specified under `define` in the .mdp file using the flag `-DPOSRES FC LIPID`. Here, a force constant of 1,000 kJ mol^-1^ nm^-2^ is chosen. 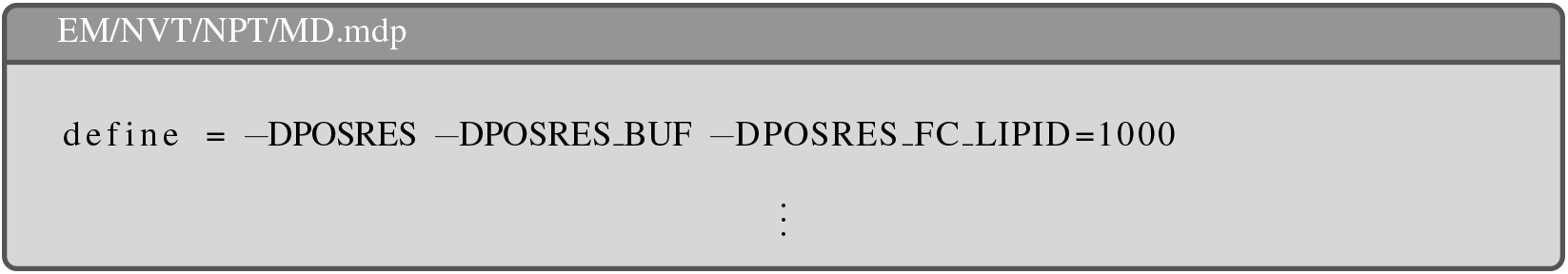
  - Subsquently, the final structure (`mc3 solvated buf cl.pdb`) can be minimized, and equilibrated with the provided parameter files: `EM.mdp`, `NVT.mdp`, and `NPT.mdp`. ~~~
gmx grompp -f EM.mdp -c phneutral.pdb -r phneutral.pdb
           -n index.ndx -p topol.top -o EM.tpr -maxwarn 2
gmx mdrun -deffnm EM -c EM.pdb -npme 0
gmx grompp -f NVT.mdp -c EM.pdb -r EM.pdb
           -n index.ndx -p topol.top -o NVT.tpr -maxwarn 1
gmx mdrun -deffnm NVT -npme 0
gmx grompp -f NPT.mdp -c NVT.gro -r NVT.gro
           -n index.ndx -p topol.top -o NPT.tpr -maxwarn 1
gmx mdrun -deffnm NPT -npme 0
~~~
  - The length of the individual runs for the thermodynamic integration is crucial for accurately estimating 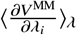 and may vary depending on the type of molecule. Proper convergence of the ensemble averages is essential, as their quality directly impacts the accuracy of subsequent constant-pH MD simulations. For the MC3 lipid, a simulation length of 600 ns per run was chosen to ensure reliable results.
  - The individual runs for the thermodynamic integration should be performed for *λ*-coordinate values between -0.1 and 1.1, with the constraint that *λ*_*i*_ +*λ*_*BUF*_ = 1.0. Therefore, a separate `MD.mdp` file must be generated for each pair (*λ*_*i*_, *λ*_*BUF*_). The script `create parameterization.py`, written by Jansen *et al*. [36], automatically modifies the `MD.mdp` file to facilitate the setup of the thermodynamic integration. **Important note:** By default, `create parameterization.py` uses a step size of 0.1 for the *λ*-coordinate. If a smaller step size is required (e.g., 0.05 as used here), the script must be modified accordingly. ~~~
python3 create_parameterization.py -f MD.mdp -r NPT.gro -c NPT.gro
                                   -n index.ndx -p topol.top
~~~
  - After the simulations are completed, the (minimum) distance between the aminolipid and the buffer particle should be checked. Fig. 3a shows all states of the aminolipid during a single window of the thermodynamic integration with the buffer particle at position `0.0 0.0 0.0`. The calculation of the minimal distance between both residues in all windows of the thermodynamic integration is shown in Fig. 3b, yielding an average minimum distance of 3.73 nm.
  - To extract the values of 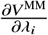 from the `ener.edr` file, use the `GROMACS` in-built function `cphmd`: ~~~
gmx cphmd -s run.tpr -e ener.edr -dvdl --coordinate no
~~~ The second column of the output file `cphmd-dvdl-1-2.xvg` contains the values of 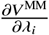 for the MC3 lipid for each frame. To allow for system equilibration, the first few nanoseconds of the simulations may be skipped. In the case of the here presented simulations, the first 10 ns were excluded from the calculation of the averages (e.g., `gmx eneconv` can be used to modify the .edr files). The polynomial coefficients were then obtained using the `fit parameterization.py` script provided by Jansen *et al*. [36]. **Note:** If `create parameterization.py` was modified to use a step size of 0.05 between the fixed *λ*_*i*_ coordinates for the creation of the .tpr files for thermodynamic integration, the `fit parameterization.py` script must also be modified to correctly find and load all required directories. For MC3, a polynomial of order 8 (i.e., 9 polynomial coefficients) was fitted according to the recommendations by Aho *et al*. [1] and Buslaev *et al*. [9]. ~~~
python3 fit_parameterization.py -f MD.mdp -m p -g DLMC -i r -fo 8
~~~ The convergence can be assessed by calculating the average values of 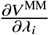 and performing the polynomial fit using only a subset of the trajectory. This subset is progressively extended by a constant value (10 ns in this case) until the entire trajectory is included. Figure 4a+b demonstrates that the averages and polynomial fits for the MC3 lipid converge after approximately 300 ns, showing minor changes thereafter.
  - The fitted polynomial coefficients found in the output file of `fit parameterization.py` next to `lambda-dynamics-group-type1-state-1-dvdl-coefficients` can be then used to update `dvdl 1` in `lambdagrouptypes.dat`. The coefficients and the charge range for the buffer sites are adapted from the paper of Buslaev *et al*. [9]. **Note:** The coefficients in the `lambdagroup-types.dat` must be written in one line, with **one** space between them. 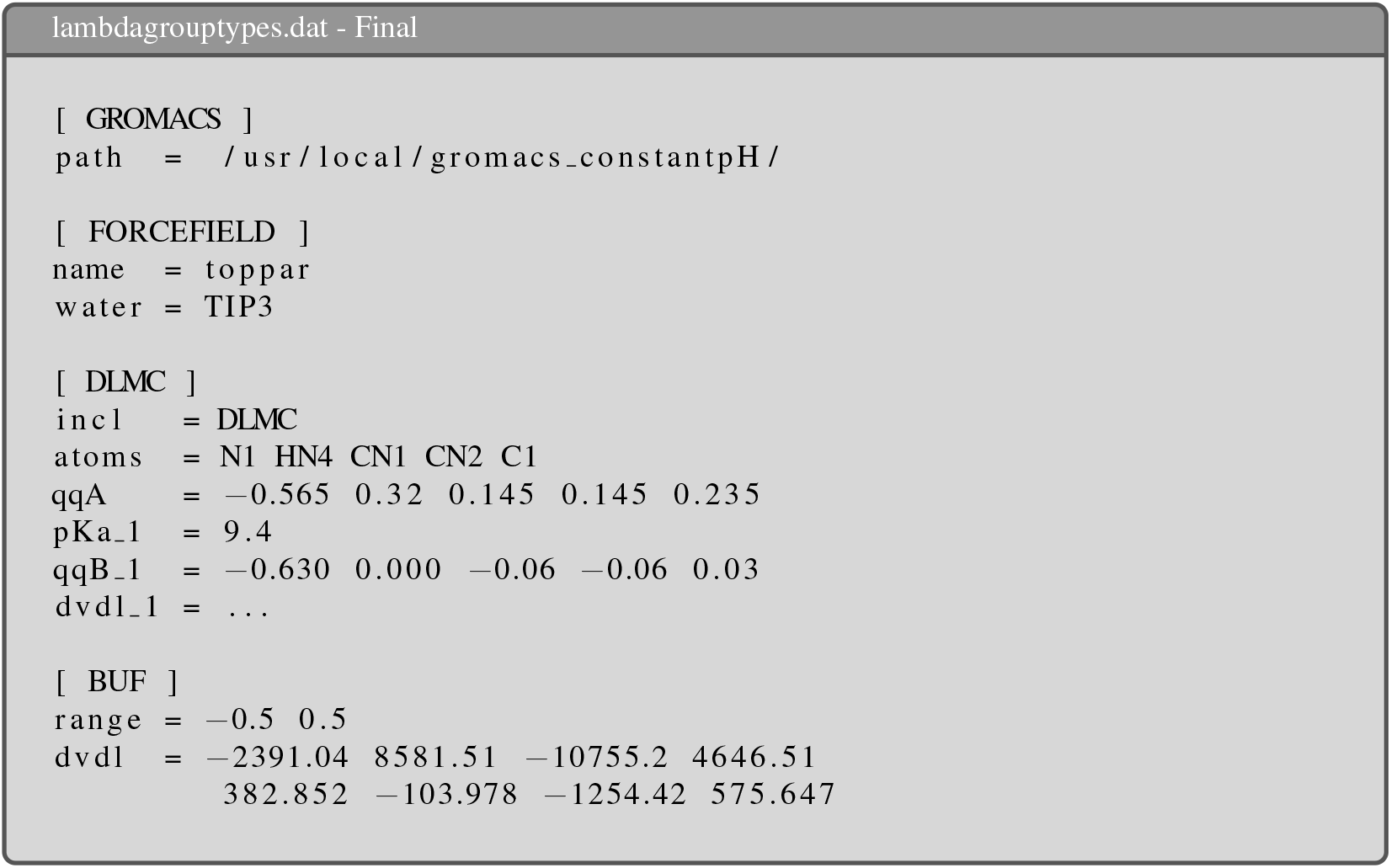
4. After the fitting routine has finished, the obtained polynomial coefficients for the MC3 lipid could be directly used for constant-pH MD simulations of complex membrane compositions. However, it is strongly recommended to validate the derived coefficients beforehand! The quality of the obtained polynomial coefficients can be assessed by evaluating their ability to cancel out the free energy of deprotonation, 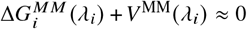,in unbiased simulations. Under ideal conditions — that is an accurate estimate of 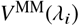 and the absence of any additional constraints on *λ*, except for *V*^bias^ (*λ*_*i*_) with a vanishing barrier height — the *λ*-coordinate is expected to have a flat distribution in the interval [0.0, 1.0] [1, 9, 36].
  - It is highly recommended to create a new, separate directory for the validation simulation and to follow the previously outlined workflow to generate a new initial structure. For the next steps, use the updated `lambdagrouptypes.dat` file with the fitted polynomial coefficients.
  - The setup of the initial structure is very similar to that used for thermodynamic integration. After solvating the system with `gmx solvate`, the `neutralize` routine from `phbuilder` is again called to add buffer particles and ions to the system: ~~~
phbuilder neutralize -f mc3_solvated.pdb -nbufs 3 -rmin 1.0 -conc 0.15 -v
                     -ignw -pname SOD -nname CLA
~~~ This command adds 3 buffer particles (`-nbufs=3`), one counter ion (i.e., chloride anion), and an ionic concentration of 150 mM Na^+^Cl^-^ to the system by replacing overlapping water molecules. The number of buffer particles allows for maximal charge fluctuation of± 1.5.
  - The .mdp files for the subsequent equilibration and final production simulations are created again with the `genparams` routine from `phbuilder`: ~~~
phbuilder genparams -f phneutral.pdb -ph 9.4 -mdp MD.mdp -dwpE 0.0
~~~ In contrast to the previous call of `genparams` for the thermodynamic integration, the `-cal` flag is not set, hence all *λ*-dependent potentials are considered during the run. To ensure that only the force field correction potential *V*^MM^(*λ*_*i*_) is effectively applied, the pH is set equal to the pK_a_ of the aminolipid, which switches off the pH-dependent potential *V*^pH^(*λ*_*i*_). Furthermore, the barrier imposed by the bias potential 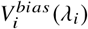 between *λ*_*i*_ = 0 and *λ*_*i*_ = 1 is set to 0.0 using `-dwpE 0.0`. Only the potential barriers at the edges remain to prevent *λ* from becoming too large (*λ*_*i*_ ≫ 1) or too small (*λ*_*i*_ ≪ 0), respectively.
  - The structure in `phneutral.pdb` can then be minimized and equilibrated in the NVT and NPT ensembles using the provided .mdp files from `genparams`. During minimization and equilibration, the positions of the buffer particles and the nitrogen atom of the MC3 lipid in this example were additionally restrained. To enhance data collection, the simulation should be repeated multiple times with different starting conditions. In the example of the MC3 lipid presented here, the equilibration (energy minimization, NVT, and NPT simulations) was performed separately for each of the 10 replica simulations.
  - The length of the unbiased simulations must be chosen to allow for a sufficient sampling of the *λ*_*i*_-coordinate. For the here presented simulations of MC3, the length of each replica simulation was set to 300 ns. Values of the *λ*_*i*_-coordinate sampled during the simulations are extracted from `ener.edr` using again `cphmd`: ~~~
gmx cphmd -s run.tpr -e ener.edr
~~~ Fig. 5a shows the probability density distributions of *λ*_*i*_ obtained from each replica simulation, along with the Boltzmann inversion of the applied bias potential *V*^bias^ (*λ*_*i*_). The distribution is approximately flat within the range of 0.0 to 1.0, indicating that the fitted *V*^MM^ (*λ*_*i*_) potential effectively cancels out the deprotonation free energy. To verify that the slight deviations do not significantly impact the results in a more realistic scenario, simulations are performed to recover the predefined *intrinsic* pK_a_ value of the aminolipid. Similar to the experimental titration method, the pK_a_ of a titratable site in constant-pH simulations can be determined by estimating the molar ratio between the protonated and deprotonated states of the residue at a given pH. However, rather than measuring this ratio across a large number of molecules at the same time (e.g., in a tube), a single aminolipid is simulated over time. The deprotonation ratio, *S*^deprot^, is calculated based on the number of simulation frames, *N*^prot^ and *N*^deprot^, where the proton is bound (*λ*_*i*_ < 0.2) or not bound (*λ*_*i*_ > 0.8), respectively [1, 9]:

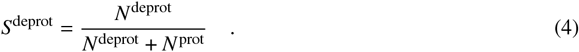

The *S*^deprot^ values across a range of pH values are related to the *intrinsic* pK_a_ of the titratable site through the Henderson-Hasselbalch equation:

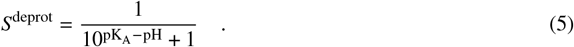
  - Similar to the first validation step, it is recommended to create a separate directory and a new initial structure for the titration simulations.
  - After inserting the required number of buffer atoms and ions with `neutralize`, ~~~
phbuilder neutralize -f mc3_solvated.pdb -nbufs 3 -rmin 1.0 -conc 0.15 -v
                     -ignw -pname SOD -nname CLA
~~~ the .mdp files can be again generated using `genparams`: ~~~
phbuilder genparams -f phneutral.pdb -ph 9.4 -mdp MD.mdp -dwpE 7.5 -titr
~~~ Adding the `-titr` flag prevents `genparams` to explicitly define a pH value in the initial MD.mdp file. Further note that the barrier between the deprotonated and the protonated state is raised to 7.5 kJ mol^-1^, which is the default value in `phbuilder`.
  - The structure in `phneutral.pdb` should be minimized and equilibrated in both the NVT and NPT ensembles using the .mdp files provided by `genparams`. During minimization and equilibration, the positions of the buffer particles and the nitrogen atom of the MC3 lipid in this example were additionally restrained.
  - The generation of .tpr files for simulations at different pH levels can be faciliated using the `create titration.py` script provided by Jansen *et al*. [36]: ~~~
python3 create_titration.py -f MD.mdp -c NPT.pdb -p topol.top
                            -n index.ndx -pH 5:15:1 -nr 10 -o r
~~~ This script prepares 10 independent replica simulations (`-nr 10`) for each pH value between 5 and 15, with a step size of 1 (`-pH 5:15:1`). Note that for the example presented here, `gen-vel = yes` and `gen-temp=310` were set in the `MD.mdp` file to vary the initial conditions of the independent replica simulations.
  - For the MC3 lipid, a simulation length of 300 ns per replica was chosen to ensure proper sampling of the *λ*-coordinate at each pH level. The values of the *λ*_*i*_-coordinate can be then again retrieved from the `GROMACS` energy output file (i.e., `ener.edr`) using `gmx cphmd`.
  - Following the approach outlined by Aho *et al*. [1], the aminolipid is considered protonated if *λ*_*i*_ < 0.2 and deprotonated if *λ*_*i*_ > 0.8 when calculating the deprotonation ratio, *S*^deprot^. Frames where *λ*_*i*_ falls within the range 0.2 ≤ *λ*_*i*_ ≤ 0.8 were excluded from the calculation of the ratio. Fig. 5b presents the values of *S*^deprot^ for the titration of the MC3 lipid used in this example, showing a sigmoidal curve characteristic of the Henderson-Hasselbalch equation.
  - A non-linear fitting routine, such as the `curve fit` function from the SciPy Python package, can be employed to determine the pK_a_ value. The following box provides an example function to calculate the pK_a_ from a set of *S*^deprot^ values, using their standard errors of the mean (i.e., the standard deviation of *S*^deprot^ across replicas) as weights for the residuals. Additionally, the standard deviation of the fitted pK_a_ value is approximated using a linear model [72]. 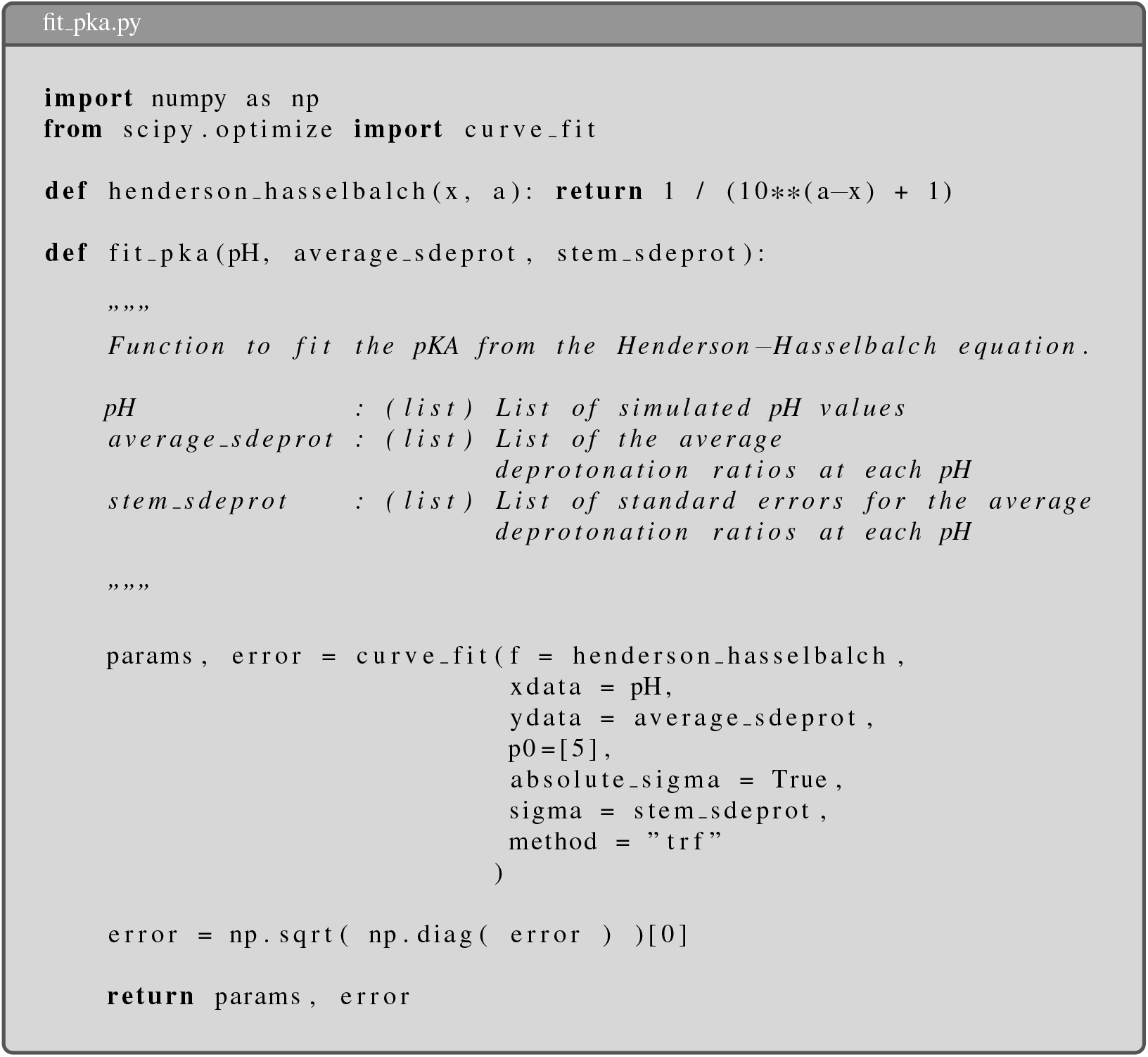
  - If the pK_a_ derived from the fitting routine matches the intrinsic pK_a_ specified in the MD.mdp file, the validation step is deemed successful. The fitted curve from the Henderson-Hasselbalch equation in Fig. 5b shows a pK_a_ of 9.39 ± 0.02, which excellently agrees with the predefined intrinsic pK_a_ of 9.4. After both validation steps have been completed successfully, the fitted polynomial coefficients can be used for further simulations. **Note:** The derived coefficients are highly sensitive to the chosen topology and force field parameters of the lipid. Any modifications to these parameters, such as adding a new functional group to the lipid headgroup, may lead to significant deviations from the validation results. Consequently, such changes would necessitate a re-estimation of the polynomial coefficients.
5. In the last validation step, the pre-defined *intrinsic* pK_a_ of MC3 could be determined from simulations of a single lipid in an aqueous phase at varying pH levels. However, previous studies [11, 29, 26, 70] have demonstrated that the pK_a_ of aminolipids — including MC3 — shifts significantly to lower values in a membrane environment. Apparently, interactions between the aminolipids and the other LNP components influences the deprotonation free energy of the titratable site. This shift has a significant influence on the potency of the LNP to deliver mRNA, and on its toxicity [51]. The following workflow guides through the simulation of a membrane system at pH 7 and how the produced data can be analyzed.

The previously generated, and shortly equilibrated membrane, is taken as an initial structure for the simulation of the MC3 lipid in a complex lipid mixture. The specific output file of the CHARMM-GUI is called `step7 1.gro` and contains the whole system, including lipids, water molecules and counter ions. However, for the constant-pH simulations enough buffer particles must be added to the system to allow for changes in the protonation state of the individual lipids. It is usually advisable to reduce the number of buffer particles in the system to reduce interactions between the titratable sites and the buffer particles [1, 9, 21]. Donnini *et al*. [21] demonstrated that, for proteins, the fluctuation in total charge is relatively small compared to the number of titratable sites, meaning that fewer buffer particles are needed to compensate for these fluctuations.

A similar assumption can be made for biomembranes containing ionizable lipids; however, the equilibration of the total charge is slower in these systems, as it may involve the reorganization of the lipid bilayer into a bi-phasic structure [69, 49]. Without knowing the *apparent* pK_a_ and the membrane structure at a specific pH beforehand, it is difficult to estimate the minimum number of buffer particles required. In this case, the number of buffer particles is set to allow for maximal charge fluctuations (i.e., 2 ·*N*^sites^ +1), with the assumption that this will not significantly affect important membrane properties. However, once the membrane structure and the number of bound protons have equilibrated, it would be possible to refine the number of buffer particles.

The following steps utilize the force field parameters provided by the CHARMM-GUI, specifically the .itp files in the `toppar` directory. For a more comprehensive force field, you can download the CHARMM36 port for GROMACS from the MacKerell Lab homepage (http://mackerell.umaryland.edu/charmm_ff.shtml, last accessed 14.11.2024).

- The number of titratable sites in the system is *N*^sites^ = 48, therefore *N*^BUF^ = 97 buffer particles are added with the `neutralize` subroutine of the `phbuilder` tool: ~~~
phbuilder neutralize -ignw -solname SOL
                     -nbufs 97 -v -p topol.top -f step7_1.gro
~~~ **Note**: CHARMM-GUI labels the water molecules as **TIP3**, since `phbuilder` expects the name **SOL**, the names in `step7 1.gro` and in `TIP3.itp` were modified accordingly.
- The input parameter file `MD.mdp` for the constant-pH simulation can be subsequently generated with `genparams`. The `step7 production.mdp` provided by the CHARMM-GUI can be used here as template, setting the simulation length to 500 ns (`nsteps=250000000`): ~~~
phbuilder genparams -f phneutral.pdb -ph 7
                    -mdp step7_production.mdp -v
~~~
- The index file produced by `genparams` contains separate groups for the atoms of each titratable group, but no groups for the membrane and the solvent needed for the temperature coupling and the center of mass removal. ~~~
gmx make_ndx -f phneutral.pdb -o index.ndx -n index.ndx
~~~ For the index groups, select `r DLMC | r DOPC | r CHL1` for the lipids and `r BUF | r CLA | r SOL` for the solvent. Name the lipid selection `MEMB` and the solvent selection `SOLV`.
- Finally, the `.tpr` can be generated using `grompp`: ~~~
gmx grompp -f MD.mdp -c phneutral.pdb -n index.ndx -o run.tpr
           -p topol.top -maxwarn 1
~~~
- The membrane structure, including the solvents, consists of approximately 70,000 atoms and can be simulated efficiently on modern GPUs using `mdrun`: ~~~
gmx mdrun -deffnm run -npme 0 -nb gpu -bonded gpu -update cpu
~~~ Specific arguments to enhance performance further depend on the available hardware. Unfortunately, the current constant-pH simulation does not allow the update and constraint steps to be shifted to the GPU.

**Fig. 2:**
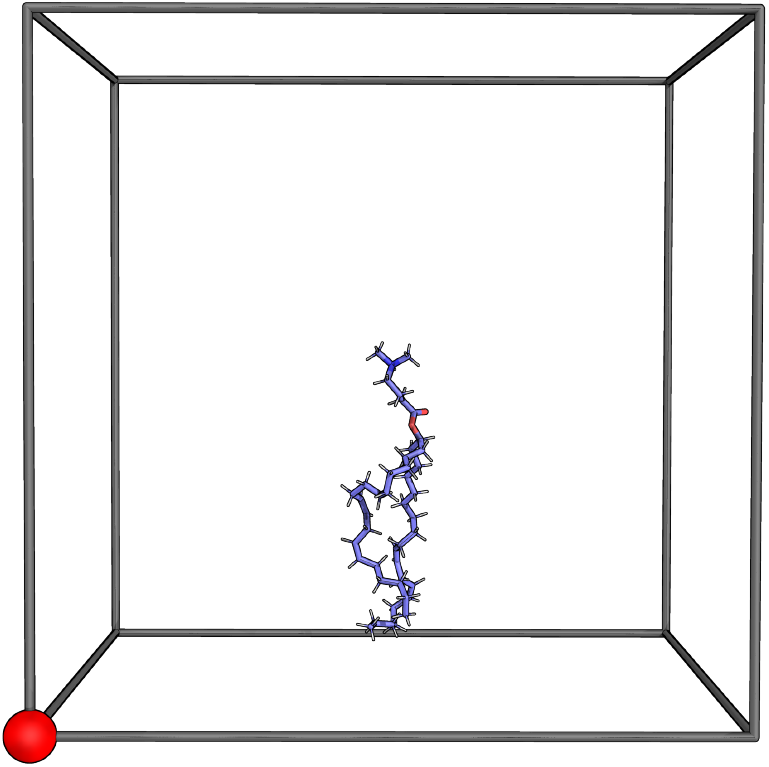
Initial structure for thermodynamic integration. The buffer particle (*red*) and the nitrogen atom in the headgroup of the MC3 lipid are fixed to their initial positions. Image created with PyMOL [60].

**Fig. 3:**
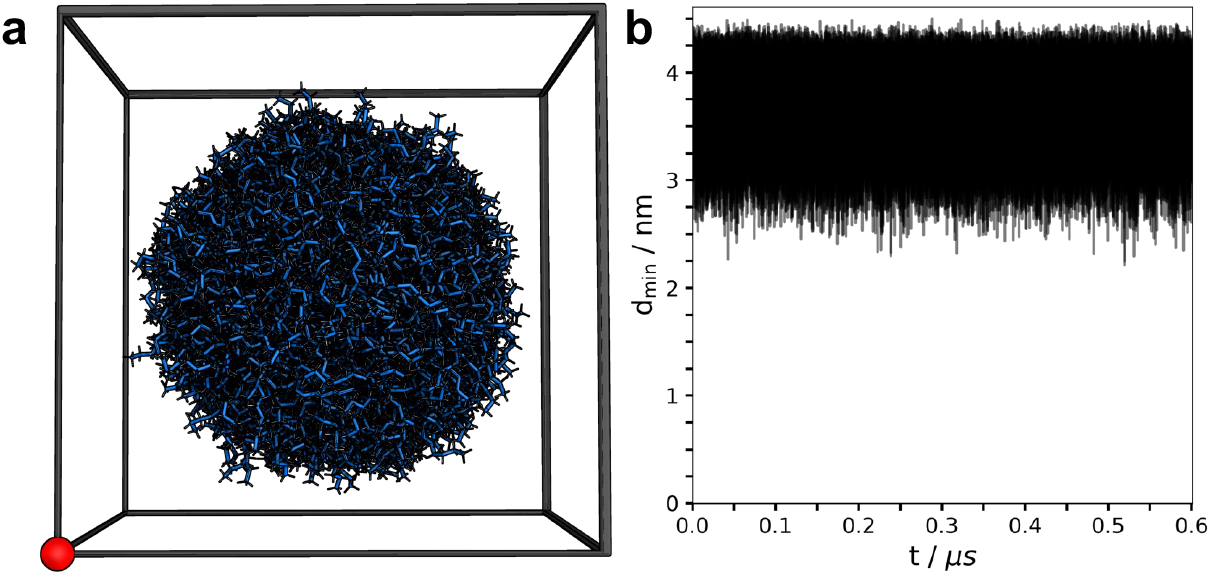
(**a**) Overlap of all positions taken by the MC3 lipid during the thermodynamic integration (shown here for *λ*_*BUF*_ = 1.10 and *λ*_*i*_ = −0.10), with the buffer particle (red) positioned at (0, 0, 0). Image was created with PyMOL [60]. (**b**) Minimum distance between the MC3 lipid and the buffer particle for all thermodynamic integration simulations. The average minimum distance between the MC3 lipid and the buffer particle is approximately 3.73 nm.

**Fig. 4:**
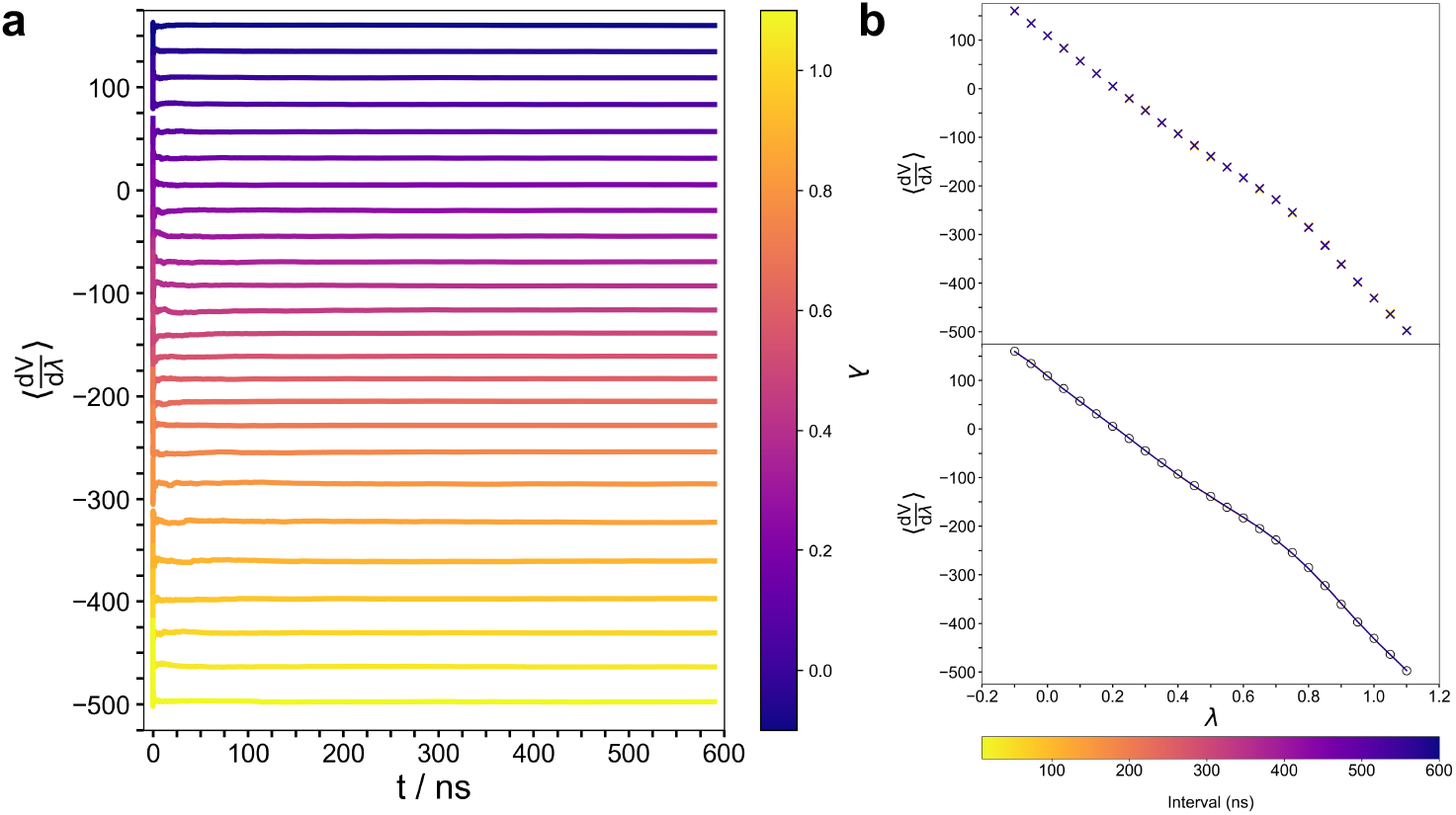
(**a**) The convergence of the gradient of *V*^MM^(*λ*_*i*_) along the *λ*_*i*_-coordinate was evaluated by progressively increasing the trajectory length used to calculate the averages, starting from 10 ns and incrementing by 1 ps. **(b)** Average gradient values of *V*^MM^(*λ*_*i*_) as a function of the *λ*-coordinate (top panel), and the evolution of the 8th-order polynomial fit for 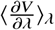 (bottom panel). Averages and fits were calculated starting from 10 ns, with the trajectory incremented in 10 ns intervals. Black circles in the bottom panel represent the final estimates for 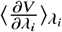, based on a total trajectory length of 590 ns.

**Fig. 5:**
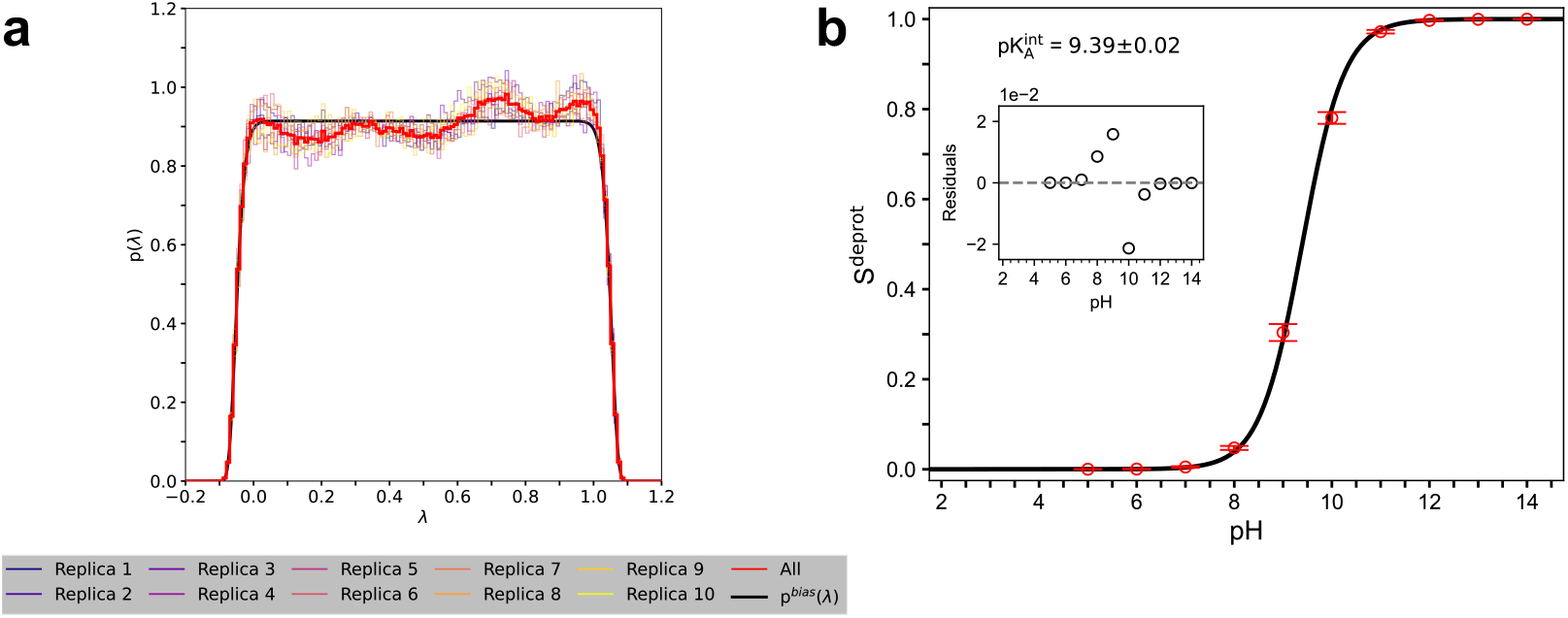
Results of the two validation steps. (**a**) Histogram of the *λ* coordinate values under V^MM^(*λ*) and V_bias_(*λ*) with a barrier height of 0.0 kJ mol^-1^. The distribution of *λ* resembles closely the probability density function dictated by *V*_bias_(*λ*), indicating successful cancellation of the deprotonation free energy by *V*^MM^(*λ*). The first 10 ns were skipped for each simulation before generating the histograms. (**b**) *S*^deprot^ of the MC3 lipid for 10 pH values. The Henderson-Hasselbalch equation was fitted to the data points (residuals of the fit are shown in the inset), weighted with the provided errors (i.e., standard deviation over 10 independent replica simulations). *N*^deprot^ and *N*^prot^ were counted over the frames of the last 290 ns.

Upon completion of the simulations, the average protonation state of the MC3 lipids and the membrane structure can be analyzed. The following section presents an example of such an analysis, focusing on key aspects of constant-pH simulations involving membranes with cationic ionizable lipids.

## 4 Results & Discussion

### Protonation States

Constant-pH simulations enable titratable sites to dynamically adjust their protonation states in response to their local environment. These interactions are primarily governed by the electrostatic potential, which must be accurately represented in the simulations. For aminolipids, the varying dielectric properties near or within the membrane are crucial factors influencing their protonation state. Inside a membrane, the protonation of an aminolipid is energetically unfavorable due to the low dielectric environment (*ϵ*_*r*≈_ 2) [10]. Conversely, protonation is more favorable in the high dielectric medium of the surrounding solvent.

Fig. 6a shows an exemplary trajectory for the nitrogen atom in the headgroup of an MC3 lipid, along with its *λ*_*i*_-coordinate at pH 7.00. When the MC3 headgroup exceeds beyond the membrane surface (e.g., between 0.46 μs and 0.49 μs), it becomes solvent-accessible, leading to a protonation of the titratable site (*λ*_*i*≈_ 0.0). Conversely, when the headgroup returns to the apolar membrane core (e.g., after ≈ 0.49 μs), the aminolipid is more likely to be deprotonated (*λ*_*i*≈_ 1.0). Altogether, the constant-pH method successfully captures the correlation between the protonation state of the titratable lipid and its local environment, consistent with the expectations based on the electrostatic potential. Notably, the application of V_bias_(*λ*) avoids extensive sampling of intermediate *λ*_*i*_ values (*λ*_*i*_ ≈ 0.5) with a relatively quick transition between the proton-bound and -unbound states.

**Fig. 6:**
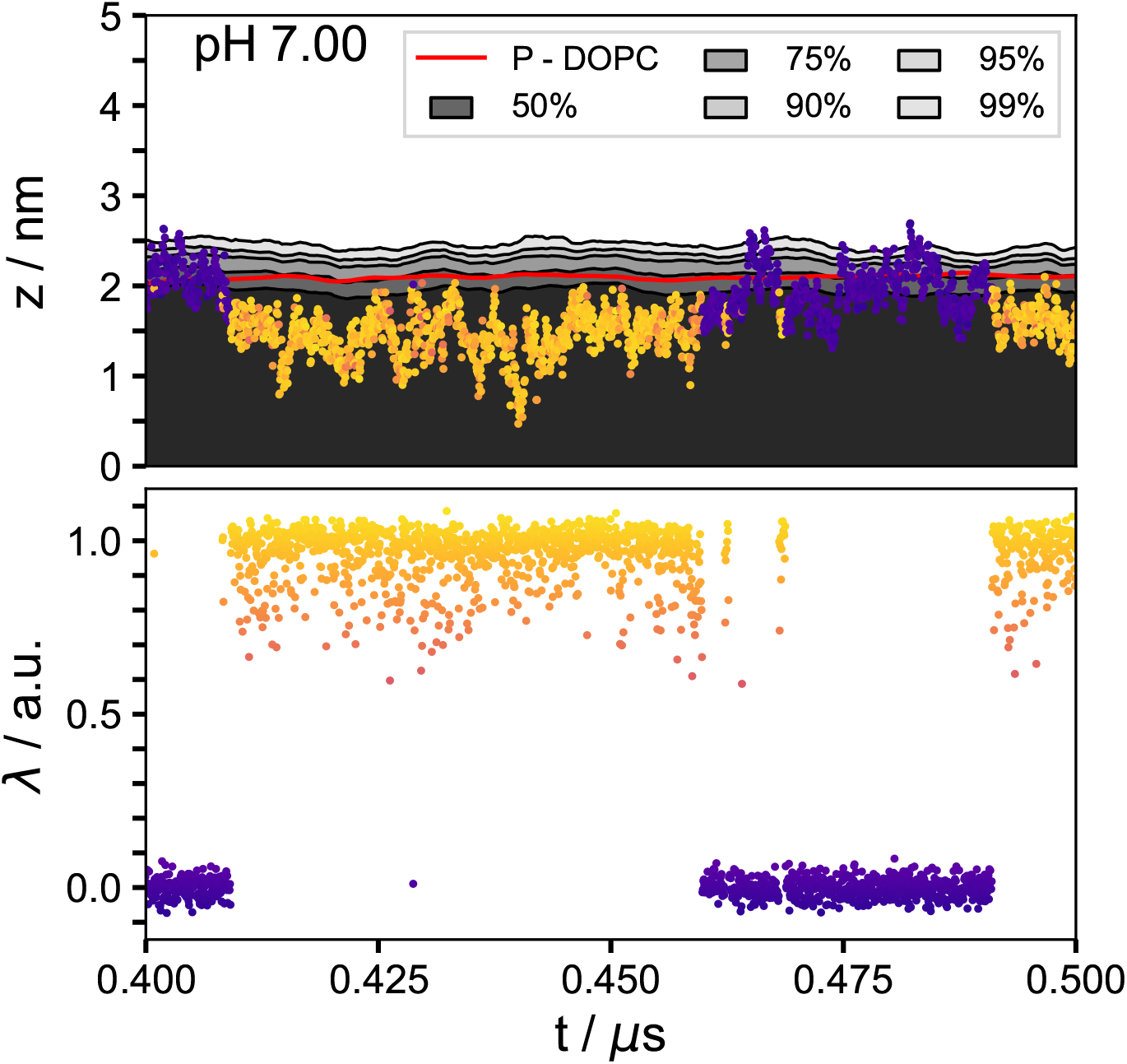
Z-position of the nitrogen atom in a single MC3 lipid, along with the median position of the phosphate group in DOPC (red line), relative to the center of the membrane. The color corresponds to the value of the *λ*_*i*_-coordinate, with blue (*λ*_*i*_ ≈ 0.0) indicating the protonated state and yellow (*λ*_*i*_ ≈ 1.0) indicating the deprotonated state. Bulk aminolipids are represented by the 50%–99% quantiles of their nitrogen z-coordinates, smoothed with a 5 ns running average. The lower panel shows the corresponding *λ*_*i*_ values.

From an intrinsic pK_a_ of 9.4, complete protonation (*S*^deprot^ ≈ 0.4 %) of MC3 in the membrane would be expected at pH 7. However, calculating the average *S*^deprot^ over the last 300 ns of the cpHMD simulation results in *S*^deprot^ ≈ 24.1 ± 1.2 %. This indicates that the lipid mixture shifts the *intrinsic* pK_a_ of the aminolipid to an *apparent* pK_a_ of approximately 7.5.

Previous studies by Carrasco *et al*. [11], Hamilton *et al*. [29], and Grava *et al*. [26] have reported even lower values for the *apparent* pK_a_ of MC3 in lipid mixtures. Measurements of the *apparent* pK_a_ in LNP formulations with molar ratios of approximately 50:10:40 MC3:DSPC:Cholesterol (including 1.5 mol% DMG-PEG2000 in [11]) using a TNS assay, *ζ* potential measurements, and umbrella sampling simulations yielded values of 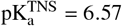 [11], 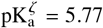 [11], and 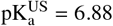 [29]. However, the latter approach does not account for phase changes in the system due to variations in environmental pH. A combined experimental/MD simulation approach to determine the protonation degree of MC3 in lipid monolayers consisting of 20:80 mol% MC3:DOPC resulted in pK_a_ = 5.73 [26].

In summary, these results highlight the substantial impact that the lipid composition has on the *apparent* pK_a_, and demonstrate that cpHMD simulations are effective in identifying these shifts, in agreement with experimental approaches.

### Structural Changes

The previous section demonstrated that constant-pH simulations effectively capture the protonation state of aminolipids and can quantify the influence of the local environment on the pK_a_. A critical follow-up question is how these protonation changes influence the overall membrane structure.

The initial membrane structure is shown in Fig. 7a, where all aminolipids are protonated (colored blue), reflecting the fixed protonation state applied during the CHARMM-GUI equilibration workflow. Over the course of constant-pH MD simulations at pH 7, a significant number of aminolipids transitioned to a deprotonated state (blue →yellow, Fig. 7a, time point 500 ns). These deprotonated aminolipids exhibit a tendency to migrate toward the bilayer core, reducing their exposure to the polar solvent. This redistribution is evident in the mass density profiles (Fig. 7b): while cholesterol and DSPC display distinct peaks at the bilayer surfaces, the MC3 lipid distribution is spread across the membrane normal (along the *z*-axis of the system).

**Fig. 7:**
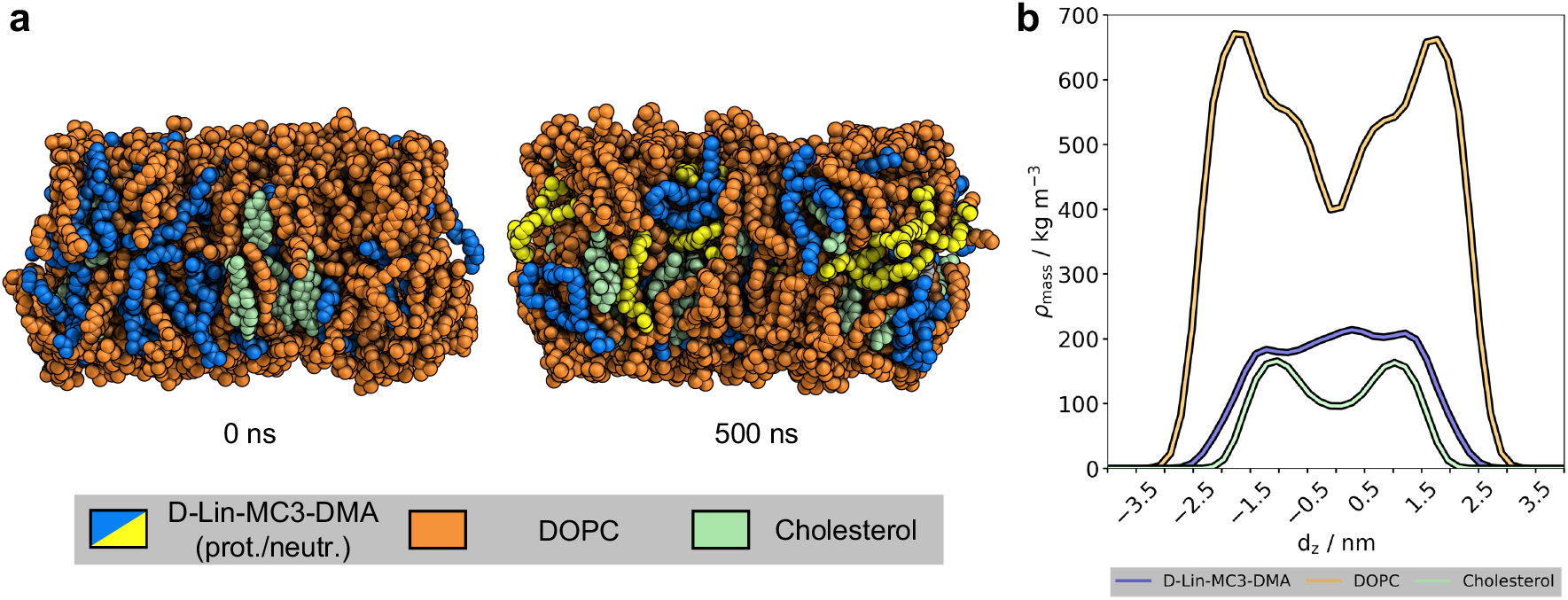
**a** Side view of the membrane structure at time points 0 ns and 500 ns during the constant-pH simulation at pH 7. Protonated aminolipids (blue) remain at the membrane surface, whereas neutral aminolipids (yellow) tend to migrate into the membrane core. Images were created with PyMOL [60]. **b** Mass density profiles of the lipids at pH 7 along the membrane normal (z-axis of the system) relative to the membrane’s center of mass. Averages were calculated over the last 400 ns.

The observed preference of uncharged MC3 lipids for deeper penetration into the hydrophobic core aligns with previous (classical) MD simulations of monolayers [26] and bilayers [34]. Similar behavior has also been reported for other aminolipids, including KC2 [56], ALC-0315 [69, 49], and SM-102 [49]. However, these earlier studies employed *static* protonation states, partly informed by experiment, lacking the dynamic protonation adjustments based on pH. It is important to note that lipid redistribution within a membrane or transitions between membrane phases upon change in pH necessitate pressure coupling in all three spatial dimensions, typically achieved using the *N PT* ensemble [71].

In summary, scalable constant-pH MD simulations provide a powerful framework for investigating complex lipid membrane systems containing hundreds of titratable aminolipids [70]. These simulations achieve high accuracy in capturing the environment-dependent protonation behavior of titratable groups while elucidating the intricate coupling between pH-dependent protonation dynamics, membrane structural changes, and lipid redistribution.

## Acknowledgements

The authors gratefully acknowledge the scientific support and HPC resources provided by the Erlangen National High Performance Computing Center (NHR@FAU) of the Friedrich-Alexander-Universität Erlangen-Nürnberg (FAU). The hardware is funded by the German Research Foundation (DFG).

